# Gene-based analysis identifies novel microRNA candidates for Autism Spectrum Disorder

**DOI:** 10.1101/2025.09.26.678002

**Authors:** Ana Rita Marques, Hugo Martiniano, Joana Vilela, Muhammad Asif, Lisete Sousa, Guiomar Oliveira, Astrid Moura Vicente

## Abstract

**Background:** Autism Spectrum Disorder (ASD) is a complex neurodevelopmental condition with unclear physiopathology. Biomarker-based diagnostic tools for early detection and targeted treatments have yet to be developed. MicroRNAs (miRNAs), which are critical regulators of genes involved in brain development and function, are emerging as promising candidates as molecular mechanisms, biomarkers, and therapeutic targets for ASD. This study aimed to identify miRNAs involved in ASD and evaluate their potential functions through *in silico* analysis.

**Methods:** A comprehensive analysis was performed to find genomic variants with a predicted functional impact on miRNA activity. Using large datasets of Single Nucleotide Variants (SNVs, *N* = 4300) and Copy Number Variants (CNVs, *N* = 3570) from ASD cases, we identified miRNAs targeted by CNVs or containing SNVs predicted to disrupt their function in ASD subjects, and compared their frequencies with controls. Selected regulatory miRNA-mRNA networks and their associated biological functions were characterized.

**Results:** Seventy mature miRNAs were identified, including both previously reported and novel ASD candidates, predicted to regulate 2742 brain-expressed genes. Enrichment analysis revealed their involvement in cellular signaling, transcription regulation, protein metabolism, and chromatin organization – biological processes strongly related to ASD. Notably, 63% of these miRNAs are predicted to target 71 known ASD risk genes and the *KCNB1, MECP2, NCKAP1* and *ZBTB20* genes are each regulated by at least four miRNAs.

**Conclusions:** This gene-based analysis identified miRNAs regulating gene networks and biological pathways implicated in brain function and plasticity, frequently disrupted in ASD. These findings strengthen the evidence for miRNA involvement in ASD, paving the way for novel diagnostic and therapeutic strategies.

## Introduction

Autism spectrum disorder (ASD) is a complex neurodevelopmental disorder with a global prevalence of 1% and a male-to-female ratio of 4.2 (1). It is characterized by deficits in social communication and interaction, along with repetitive behaviors and interests, often co-occurring with other comorbidities that affect patient’s daily activities (2). Strong evidence suggests that ASD is highly heritable, with genetic factors accounting for 50 to 80% of the familial ASD risk (3,4). Many different genes and genomic regions have been associated with ASD, highlighting convergent biological processes such as gene regulation, chromatin remodeling, and neuronal communication (5–9). However, variants in protein-coding genes explain only a proportion of ASD cases (8–11) and, although changes in gene expression have been well documented in ASD (12) little is known concerning the contribution of regulatory mechanisms to the disorder. This emphasizes the need to explore the role of regulatory sequences outside protein-coding regions, and in particular the small noncoding RNA (sRNA) molecules that control gene expression at the post-transcriptional level.

MicroRNAs (miRNAs) are small noncoding RNAs, 18–25 nucleotides in length, that complementarily bind target messenger RNAs (mRNAs) and regulate the expression of >60% of the transcriptome. They derive from hairpin precursor transcripts and are mainly implicated in silencing of gene expression, translational repression, and mRNA degradation (13,14), but have also been associated with upregulated gene expression (15). A single microRNA can regulate different mRNAs and one mRNA can be targeted by multiple microRNAs. This way, miRNAs–mRNAs form complex gene regulatory networks that participate in various biological processes in the brain, such as synaptic plasticity, neurogenesis, and neuronal maturation (16). Accordingly, miRNAs have important functions during brain development and throughout life, and their dysregulation contributes to human pathologies, including ASD, Attention-Deficit/Hyperactivity Disorder (ADHD), and Schizophrenia (17–19). Importantly, independent studies have demonstrated that miRNA expression profiles are altered in several tissues of ASD patients, including blood, saliva, and brain (17,20–22), with specific miRNAs consistently reported in 4 or more studies (miR-155-5p, miR-146a-5p, and miR-106a-5p) (17,20–28). These alterations suggest that miRNAs are involved in ASD pathophysiology. While most of the studies so far have been focused on miRNA expression, a few have explored the impact of genomic single nucleotide variants (SNVs) in miRNA genes (29–31) in small datasets (29,31), with intriguing findings that beseech more extensive investigation.

In this study, we aimed to investigate the potential contribution of regulatory variants to ASD risk, through the identification of genomic variants that have a predicted functional impact on miRNA functions in individuals diagnosed with ASD. For that, we performed a gene-based analysis using large datasets containing information on SNVs and Copy Number Variants (CNVs) of ASD patients and controls. We placed a strong emphasis on crucial features of miRNAs, in particular the highly conserved seed domains in the mature sequence with perfect complementarity between a miRNA and its target mRNA, which are essential for target-binding recognition. Additionally, the well-defined structural and primary sequence features of hairpin miRNAs such as CNNC, basal-UG and apical UGU/UGUG motifs are essential for miRNA functions and were well considered (14). Both SNVs and structural variants, namely CNVs, affecting any of these structures can interfere with the expression of mature miRNAs and influence miRNA targeting ability (32–34), and are worthy of more extensive exploration. To further understand the biological context of changes in miRNA genes, we used functional enrichment analysis to identify miRNAs-mRNAs interaction networks involved in biological pathways and processes implicated in ASD.

## Methods

The workflow used in this study for identifying miRNA variants and associated biological processes relevant for ASD is represented in Figure 1.

**Figure 1.**
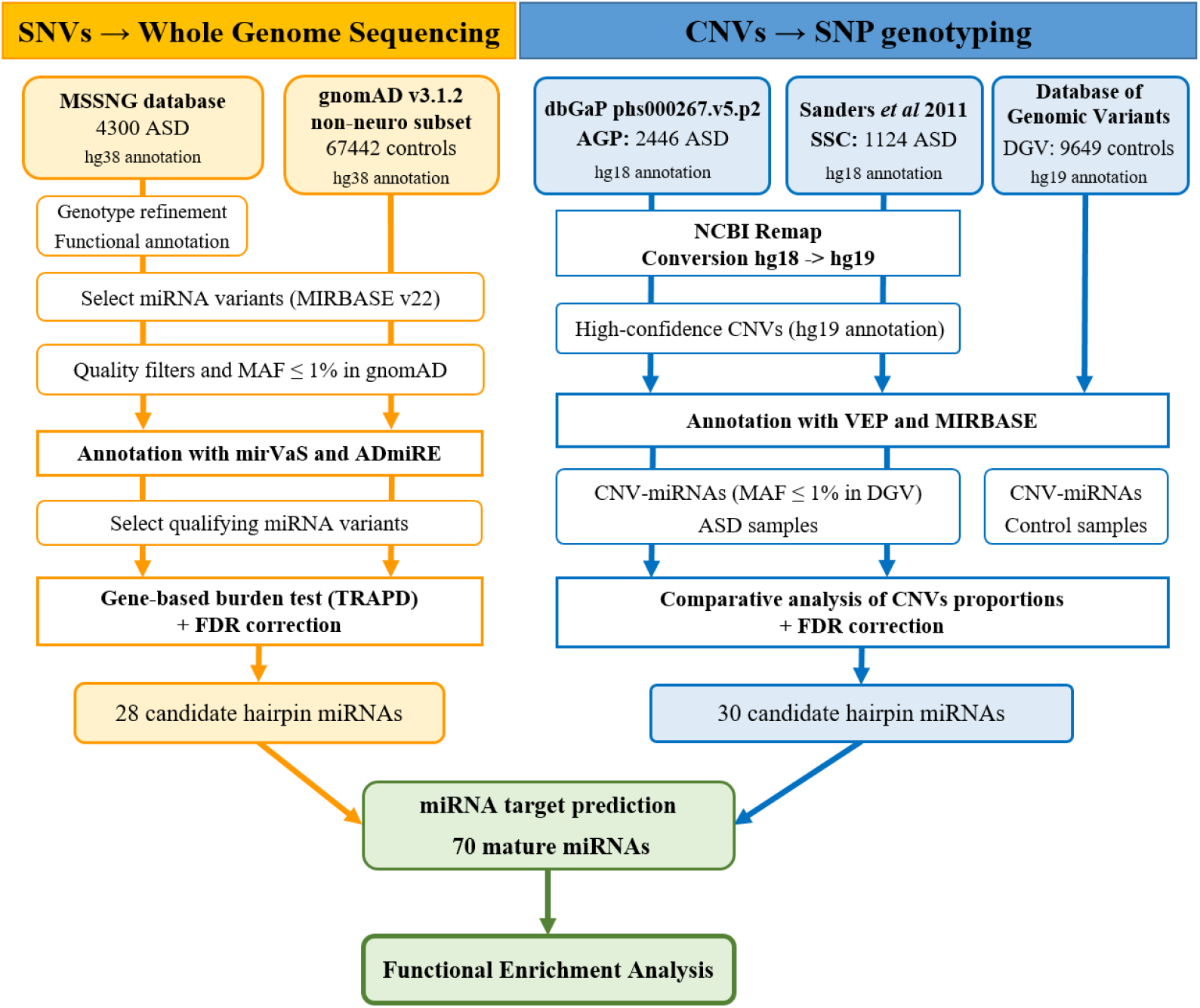
Workflow for identifying miRNA variants and associated biological processes in ASD pathophysiology. Large population datasets were analyzed to identify SNVs (orange) and CNVs (blue) targeting miRNA genes in individuals with ASD. Target genes of candidate miRNAs were predicted followed by enrichment analysis. Legend: SNVs, Single Nucleotide Variants; CNVs, Copy Number Variants; MSSNG, Autism Speaks database; gnomAD, Genome Aggregation Database; AGP, Autism Genome Project; SSC, Simons Simplex Collection; DGV, Database of Genomic Variant; miRNA, microRNA; MIRBASE, the microRNA database; MAF, Minor Allele Frequency; mirVaS, tool that predicts the impact of genetic variants on miRNAs; ADmiRE, Annotative Database of miRNA Elements; TRAPD, Testing Rare vAriants using Public Data; FDR, False Discovery Rate; VEP, Variant Effect Predictor.

### Genomic datasets

The SNV dataset included Whole-Genome Sequencing (WGS) data from 4300 ASD subjects from the MSSNG database (MSSNG, https://research.mss.ng/) (9,11). A total of 3570 ASD subjects with CNV data were analyzed from the Autism Genome Project consortium (AGP, *N* = 2446) (6) and from the Simons Simplex Collection (SSC, *N* = 1124) (5) datasets. Individuals with ASD from these studies met the criteria for autism or ASD on the ADOS and ADI-R diagnostic measures or were diagnosed by an expert clinician according to the DSM (IV or 5 editions) (5,6,9,11). As reference control population, WGS data from 67442 unrelated individuals without a neurological condition were obtained from the non-neuro subset Genome Aggregation Database v3.1.2 (gnomAD, https://gnomad.broadinstitute.org/downloads) (35) and CNVs from 9649 unrelated individuals with no history of neuropsychiatric disorder were obtained from the Database of Genomic Variant (DGV: http://dgv.tcag.ca/dgv/app/home) (36).

### Variant Analysis and Prioritization

For SNV analysis, WGS and variant detection for both cases and controls were performed as previously described (9,11). Only variants located on miRNA precursor stem-loop sequences with a Minor Allele Frequency (MAF) ≤ 1% or not detected in gnomAD (v3.1.2) were considered. To predict the functional and structural impact of the identified miRNA variants, they were annotated with mirVaS (37) and the Annotative Database of miRNA Elements (ADmiRE) (38). Variants were predicted to have damaging effects if located on seed, mature, basal and apical motifs of the miRNA sequence, or predicted to change the structure of the hairpin. Only these were selected from both ASD and control datasets and are referred to as “qualifying variants”.

CNV discovery was previously performed using Illumina SNP genotyping data as previously described (5,6,39,40). For this study, the genomic content of CNVs was remapped from hg18 to hg19 and re-annotated. The rare CNVs overlapping miRNAs (CNV-miRNAs) with a frequency <1% in controls were identified and considered for gene-based analysis.

### Gene-based analysis to select miRNA candidates

Two different gene-based analysis methods were used to identify candidate miRNAs from SNV and CNV datasets.

For SNVs, the software package TRAPD (Testing Rare vAriants using Public Data) was used to perform gene-based burden testing using individual-level genotyping data from ASD cases (MSSNG) and summary-level public control data (gnomAD), as described by Guo *et al* (41). A one-sided Fisher’s exact test was applied to determine if cases have a higher burden of qualifying variants than controls for each gene, under both a dominant and a recessive model. Burden testing was performed using the “burden_test.R” function in TRAPD (available on https://github.com/mhguo1/TRAPD). Using the “p.adjust” function, *p*-values were adjusted for multiple testing using False Discovery Rate (FDR) correction at α = 0.05.

For CNVs, a two proportions comparison test was applied to establish whether the proportion of ASD-subjects carrying CNVs targeting a given miRNA gene (*p*_*ASDcases*_ ) is higher than the proportion of control subjects carrying CNVs targeting that same miRNA gene (*p*_*controls*_), where *p* stands for proportion. This analysis consists of a one-tailed test for which the null hypothesis (H0) is defined by *p*_*ASDcases*_ ≤ *p*_*controls*_, and the alternative hypothesis (H1) is given by *p*_*ASDcases*_ > *p*_*controls*_. FDR correction for multiple testing was applied, with significance at α = 0.05. All statistical analysis were performed using R.3.2.3 software.

### Identification of miRNA target genes

The miRNA targets were retrieved from three miRNA target prediction databases: mirWalk v3.0 (updated January 2022) (42), miRDB v6.0 (updated June 2019) (43) and miRTarBase v9.0 (updated January 2022) (44). We used a conservative strategy to analyze the predicted target genes by restricting the analysis to the targets that were predicted *in silico* in both mirWalk (score ≥ 0.95) and miRDB (score ≥ 80) and/or experimentally validated in miRTarBase. The predicted miRNA targets were converted to the official gene symbol that has been approved by the HUGO Gene Nomenclature Committee (HGNC) (45), using the HGNC BioMart server (data retrieved on August 16th, 2022).

### Functional analysis of miRNA targets

The functional impact of candidate miRNAs was assessed through the enrichment analysis of the miRNA target genes using the g:Profiler tool (version e108_eg55_p17_0254fbf, retrieved on March 8th, 2023), with FDR correction for multiple testing (α = 0.05) (46). The data retrieved from g:Profiler was derived from the Reactome, KEGG and Gene Ontology (GO) databases. To increase the specificity of enrichment results, the more representative pathways from the manually curated Reactome database (47) were chosen by grouping top level pathways according to Reactome hierarchy. To identify more specific GO terms, the terms with more than 600 genes within the group were excluded, and the remaining redundant GO terms were removed as described previously (48). To evaluate potential interactions between the miRNA targets, a protein-protein interaction (PPI) network was created using the STRING database (v11.5, https://string-db.org/), with species limited to “*Homo sapiens*” and confidence score ≥ 0.7. The resulting network map comprising the known and predicted PPIs and miRNA-mRNA regulatory interactions was visualized using Cytoscape (version 3.10.2) (49).

### Biological relevance of miRNAs and target genes

Information on ASD risk genes and brain expression was used to assess the biological relevance of the identified miRNAs. Overlap between miRNA target genes and 1220 ASD candidate genes from the manually curated SFARI Gene database (released on 13 January 2025, https://gene.sfari.org/) was analyzed. In this database, genes are ranked into four categories: syndromic, high confidence, strong candidates and suggestive evidence. The expression of miRNAs and their target genes was assessed in the adult human brain and during early stages of brain development, using data retrieved from GTEx portal (50), Human Protein Atlas (HPA) (51), DIANA-mITED (52), BrainSpan Atlas (53) and EMBL-EBI Expression Atlas (54).

## Results

Figure 1 illustrates the overall approach for discovering miRNA gene variants and identifying biological processes relevant to ASD pathophysiology.

### Discovery of SNVs in microRNA genes

MSSNG genome-sequencing data from 4300 ASD individuals was analyzed to identify SNVs in miRNA genes. A total of 19819 SNVs were detected in 93% (1784 out of 1913) of the human microRNA genes annotated in MIRBASE v22, providing near-complete coverage of all known miRNA genes.

SNVs located in functionally relevant regions of the miRNA hairpin, including the seed, mature and motif sequences, (see Figure 2A) were prioritized as qualifying variants, due to their potential to disrupt miRNA processing, stability, or target recognition. In total, 3660 qualifying variants in 1378 miRNA genes were identified (Supplementary Table 1). Importantly, a subset of these miRNAs had previously been implicated in ASD: 66 miRNAs were described with altered expression in ASD patients when compared to controls in at least two independent studies (17,20–23,25–28,55–83), while 58 miRNAs were associated with ASD in genetic studies (29–31,84–87) (Supplementary Table 2).

**Figure 2.**
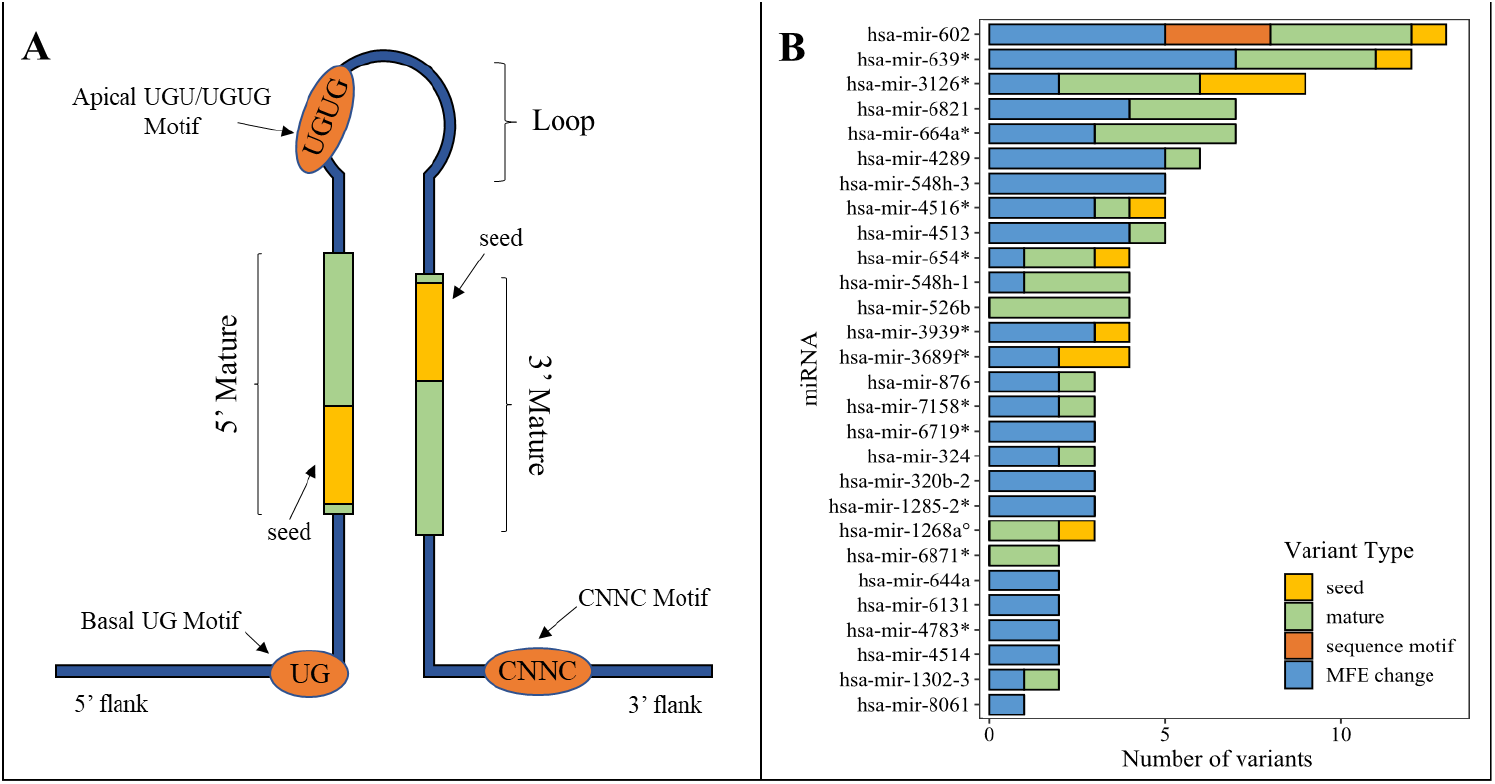
Type of qualifying variants identified in candidate miRNAs. (A) Schematic representation of stem-loop structure with highlighted distinct functional miRNA gene regions (adapted from ADmiRE (38) [Oak 2019). (B) Number of different SNVs identified in the 28 miRNAs enriched in ASD subjects (adjusted *p*-value < 0.05). Legend: *recessive model; °both dominant and recessive models; MFE change, predicted to change the structure of the hairpin according to the minimal free energy (MFE) structure.

### Gene-based analysis identifies miRNA candidates

To assess the impact of miRNA qualifying variants at the gene level, a gene-based burden test was performed to compare the number of individuals carrying such variants in each miRNA gene between ASD cases (*N* = 4300) and gnomAD non-neuro controls (*N* = 67442). As shown in Figure 2B, the burden test revealed that variants in 28 miRNAs genes were significantly enriched in ASD patients, with 15 enriched in the dominant model, 12 enriched in the recessive model and one enriched in both models (Supplementary figure 1 and supplementary table 3). Most of these 28 miRNAs are expressed in brain and/or during neurodevelopment, and 21 are novel for ASD. Seven of the 28 miRNAs, namely the hsa-mir-324, hsa-mir-654, hsa-mir-664a, hsa-mir-876, hsa-mir-1268a, hsa-mir-4516 and hsa-mir-6821 were previously reported in CNVs and/or gene expression studies of ASD patients (21,24,56,57,74,77,84,87,88). As shown in Figure 2B, the hsa-mir-602 contained the highest number of qualifying variants (*n* = 13), identified in 3% (137/4300) of the ASD cases, followed by hsa-mir-639 and hsa-mir-3126 with 12 and 9 qualifying variants identified in 1% (43/4300) and 1.3% (56/4300) of the ASD cases, respectively. Interestingly, two significantly enriched miRNA genes, hsa-mir-548h-3 and hsa-mir-548h-1, contained 9 qualifying variants identified in 2.9% (125/4300) of the ASD cases, and they give rise to the same mature sequence that is expressed in brain and during neurodevelopment – the hsa-miR-548h-5p.

### Discovery of microRNA genes targeted by CNVs

A total of 2350 CNVs targeting miRNA genes were identified in AGP (n = 2026, supplementary Table 4) and SSC (n = 324, supplementary Table 5) ASD samples. Eighteen miRNA genes were targeted by CNVs exclusively in ASD patients, but in none of the 9649 controls, with a frequency > 0.01% (corresponding to at least 4 ASD cases in the AGP and SSC datasets). From these, 12 miRNA genes were found in duplicated CNVs, 4 were found in deleted CNVs, and 2 were identified both in duplicated and deleted CNVs, as shown in Table 1. There were 6 miRNA genes (hsa-mir-4436b-2, hsa-mir-3179-3, hsa-mir-3180-3, hsa-mir-3680-2, hsa-mir-484 and hsa-mir-4767) that were targeted by CNVs exclusively in ASD patients from the AGP and SSC datasets and none of the controls.

**Table 1.** Frequencies observed for miRNA genes targeted by CNVs exclusively in individuals with ASD from the AGP (*N* = 2446) and/or SSC (*N* = 1224) datasets, when compared to controls from the DGV dataset (*N* = 9649).

| MIRBASE ID<br>(stem-loop) | Cytoband | CNV type | AGP $n$ (%) | SSC $n$ (%) | SSC and AGP<br>Total $n$ (%) |
| --- | --- | --- | --- | --- | --- |
| hsa-mir-11399 | 1p13.2 | gain | 4 (0.16) | 0 | 4 (0.11) |
| hsa-mir-1302-2 | 1p36.33 | gain | 7 (0.29) | 0 | 7 (0.19) |
| **<u>hsa-mir-4436b-2</u>** | 2q13 | gain | 14 (0.57) | 1 (0.08) | 15 (0.41) |
| **<u>hsa-mir-4436b-2</u>** | 2q13 | loss | 11 (0.45) | 3 (0.25) | 14 (0.38) |
| hsa-mir-10525 | 7q11.23 | gain | 0 | 4 (0.33) | 4 (0.11) |
| hsa-mir-1299 | 9p11.2 | gain | 12 (0.49) | 0 | 12 (0.33) |
| hsa-mir-4477b | 9q13 | loss | 6 (0.25) | 0 | 6 (0.16) |
| hsa-mir-4511 | 15q22.31 | loss | 4 (0.16) | 0 | 4 (0.11) |
| hsa-mir-3179-1 | 16p13.11 | gain | 4 (0.16) | 0 | 4 (0.11) |
| **<u>hsa-mir-3179-3</u>** | 16p12.3 | gain | 2 (0.08) | 2 (0.16) | 4 (0.11) |
| hsa-mir-3179-4 | 16p12.3 | gain | 3 (0.12) | 0 | 3 (0.08) |
| hsa-mir-3180-1 | 16p13.11 | gain | 4 (0.16) | 0 | 4 (0.11) |
| **<u>hsa-mir-3180-3</u>** | 16p12.3 | gain | 2 (0.08) | 2 (0.16) | 4 (0.11) |
| **<u>hsa-mir-3680-2</u>** | 16p11.2 | gain | 3 (0.12) | 3 (0.26) | 6 (0.16) |
| **<u>hsa-mir-3680-2</u>** | 16p11.2 | loss | 3 (0.12) | 6 (0.49) | 9 (0.25) |
| **<u>hsa-mir-484</u>** | 16p13.11 | loss | 3 (0.12) | 1 (0.08) | 4 (0.11) |
| hsa-mir-6511b-2 | 16p13.11 | gain | 4 (0.16) | 0 | 4 (0.11) |
| hsa-mir-6770-1 | 16p13.11 | loss | 4 (0.16) | 1 (0.08) | 5 (0.14) |
| hsa-mir-6770-3 | 16p12.3 | gain | 2 (0.08) | 2 (0.16) | 4 (0.11) |
| **<u>hsa-mir-4767</u>** | Xp22.31 | gain | 11 (0.45) | 6 (0.49) | 17 (0.46) |

The table shows genes observed exclusively in both the AGP and SSC datasets (in bold and underlined), or exclusively only in one of the datasets (not in bold). CNVs, Copy Number Variants; AGP, Autism Genome Project; SSC, Simons Simplex Collection.

A two-proportion comparison test was performed separately for each of the two independent ASD datasets to identify miRNAs genes more frequently targeted by CNVs in cases. As shown in Table 2, the analysis revealed 13 miRNA genes targeted by a higher proportion of duplicated CNVs in ASD subjects (11 in the AGP and 4 in the SCC datasets). Notably, 2 miRNA genes (hsa-mir-4771-1 and hsamir-548x) showed a higher proportion of duplicated CNVs in ASD subjects from both the AGP and SCC datasets.

**Table 2.**
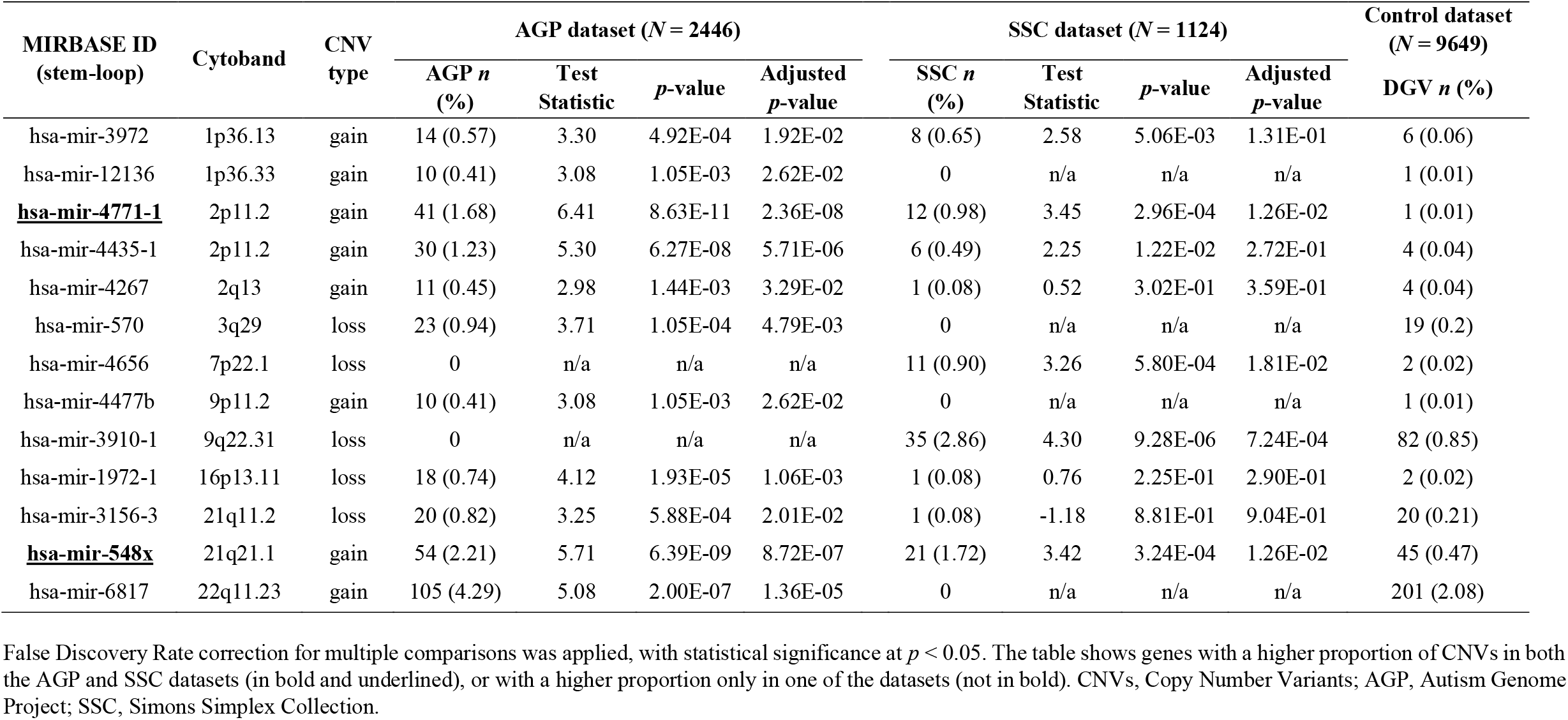
Summary statistics for the miRNA genes with a higher proportion of CNVs in individuals with ASD from AGP and datasets when compared to controls from the DGV dataset.

Overall, 30 miRNAs genes targeted by CNVs were identified in ASD patients, from which 22 are novel findings in ASD. The remaining 8 were previously associated with ASD in CNV and gene expression studies (17,22,23,57,68,77,84,85,89). Interestingly, some of the miRNA genes identified by this CNV analysis are located in regions that have previously been strongly associated with ASD risk, and may be contributing to the ASD phenotype (3q29, 7q11.23, 16p11.2, 16p13.11 and 16p13.3, Supplementary Figure 2).

### Analysis of miRNA target genes

With this analysis, 58 miRNA genes were identified as ASD candidates (28 SNVs and 30 CNVs). Because one or two mature miRNAs can be processed from each stem-loop structure, these 58 hairpins corresponded to 70 mature miRNAs (Supplementary Table 6). For these 70 miRNAs, 2841 experimentally validated and predicted mRNA targets were identified, resulting in 3773 miRNA-mRNA interactions (supplementary Table 7). Of these interactions, 97% (3671/3773) involved 2742 target genes that are expressed in the adult brain and 93% (3526/3773) involved 2632 target genes expressed during early brain development. Interestingly, 425 of the miRNA target genes have an elevated expression in the brain compared to other tissue types, suggesting they have an important role in brain function.

Further analysis evaluated whether these target genes had been previously implicated in ASD. Among the 2742 brain-expressed miRNA targets, 71 were classified as high-confidence ASD candidate genes in the SFARI database. These genes are regulated by 63% (44/70) of the mature miRNAs identified in this study, supporting a potential role for these miRNAs in ASD. An integrative network illustrating these miRNA-mRNA interactions and the PPI of the miRNA targets is shown in Figure 3. Twenty-one genes are targeted by more than one miRNA, with four genes (*KCNB1, MECP2, NCKAP1* and *ZBTB20*) being targeted by at least four miRNAs. As shown in Figure 3 (red arrow), seven of these miRNAs (hsamiR-6817-3p, hsa-miR-3680-3p, hsa-miR-4516, hsa-miR-484, hsa-miR-526b-3p, hsa-miR-1299, and hsa-mir548x-3p) target 5 to 9 genes listed as ASD candidates in SFARI category 1.

**Figure 3.**
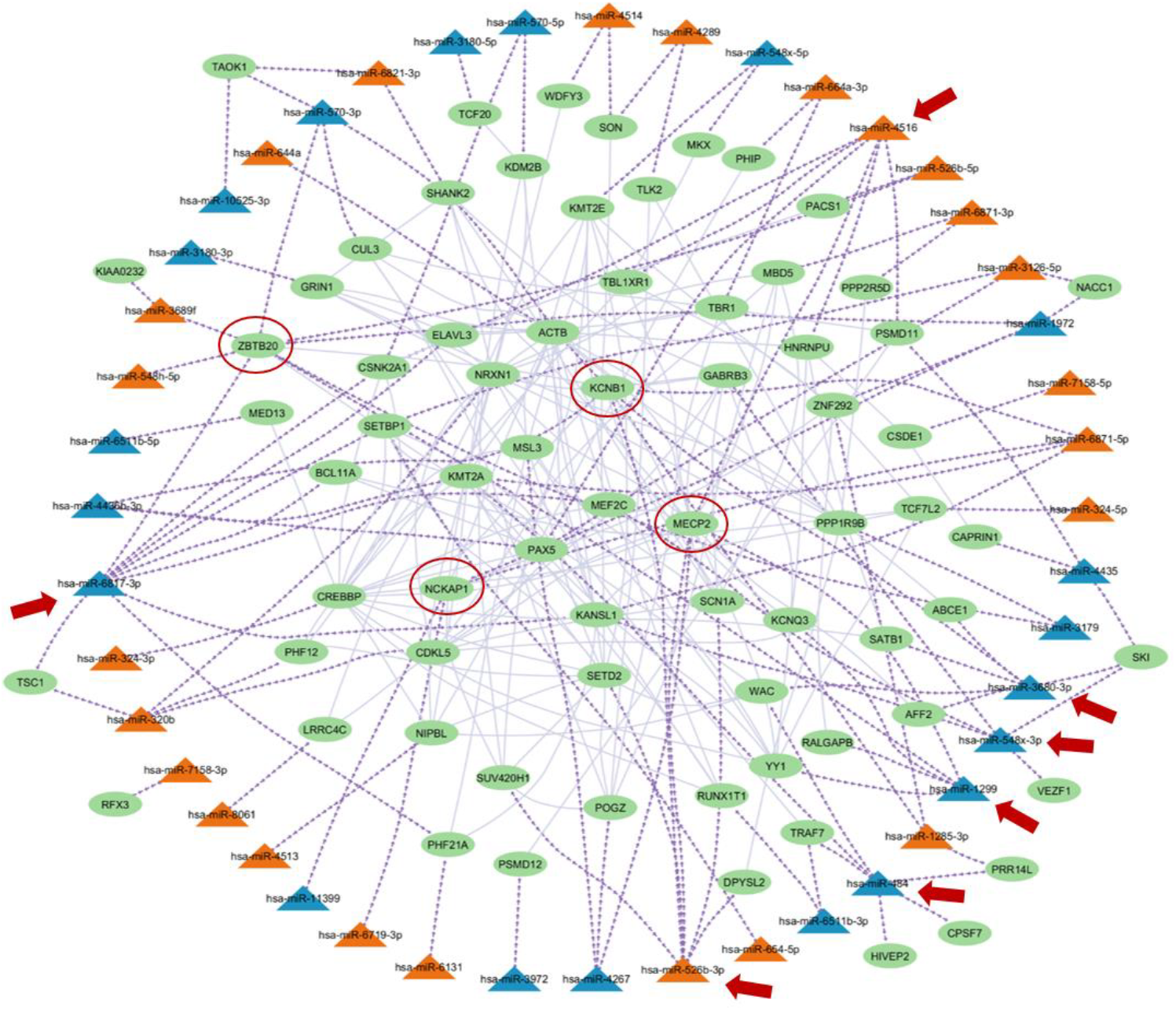
Network diagram of miRNA-mRNA and PPI interactions for miRNA targets that are high confidence ASD candidates in SFARI database. Node shape represents genes (circle) and miRNAs (triangle), edges shape represent miRNA-mRNA (dashed/arrow purple) and PPI (straight grey) interactions. Node fill color represent genes (green) from Sfari category 1 that are targeted by candidate miRNAs and candidate miRNAs observed in SNV (orange) and CNV (blue) analysis. Red arrow represents the miRNAs that target more than 5 ASD candidate genes. Red circle highlights genes that are target by at least 4 miRNAs.

### Functional enrichment analysis

To explore the functional relevance of the 70 mature miRNAs, an enrichment analysis of their 2742 brain-expressed targets was conducted (supplementary Table 8). Reactome pathway analysis identified a significant overrepresentation of pathways involved in cellular signaling (including the MAPK, PI3K-Akt, Neurotrophin and RHO GTPases signaling pathways), transcription regulation, protein metabolism and chromatin organization (Figure 4A). GO analysis revealed significant enrichment (adjusted *p*-value < 0.001) for molecular functions related to transcription regulation and protein binding (Figure 4B), and for biological processes associated with neuron differentiation, translation, histone modification, and cell signaling (Figure 4C). Cellular component analysis showed significant enrichment (adjusted *p*-value < 0.001) in synaptic structures and regulatory complexes, including the presynapse, postsynapse, neuron-to-neuron synapse, glutamatergic synapse, transcription regulator complex, transcription repressor complex, and histone acetyltransferase complex (Figure 4D). These findings highlight the involvement of these genes in critical mechanisms underlying brain function and neurodevelopment, for which there is previous strong evidence of being compromised in ASD.

**Figure 4.**
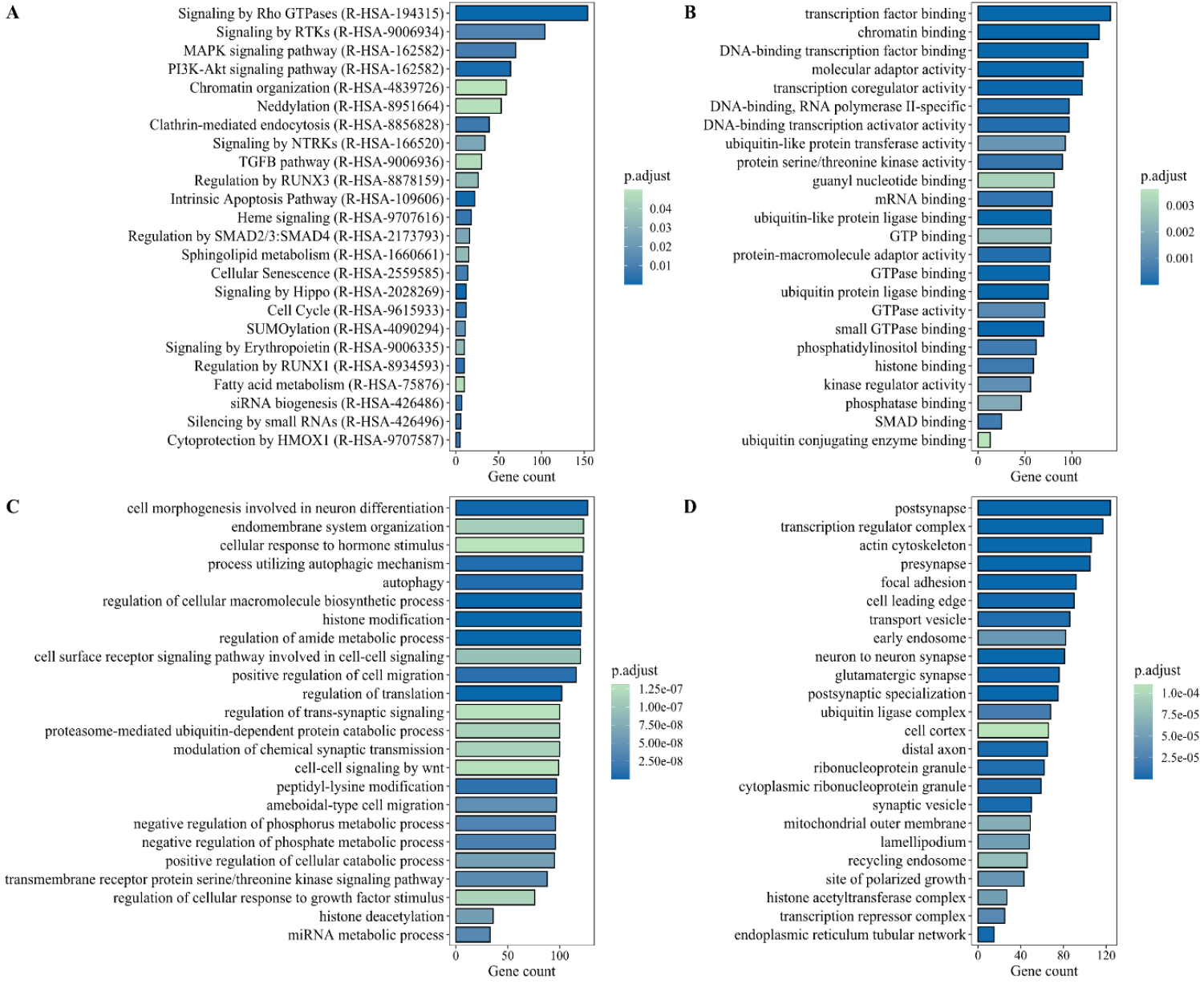
Functional enrichment analysis of the 2745 miRNA targets. **(A)** Results for the Reactome pathways; **(B)** Results for GO Molecular Function; **(C)** Results for GO Biological Process; **(D)** Results for GO Cellular Component. P.adjust is the p-value after the FDR correction and smaller values indicate the enrichment is more significant. GO is Gene Ontology.

The specific role of each miRNA in these mechanisms was also investigated. Notably, 91% (64/70) of the identified mature miRNAs were involved in cell signaling, including 55 in the Rho GTPase, 46 in the PI3K-Akt, and 47 in the MAPK signaling pathways. In addition, 60% (42/70) of these mature miRNAs were also involved in chromatin organization, 59% (41/70) in gene expression, and 43% (39/70) in protein metabolism (Figure 5). Overall, our analysis suggests that these miRNA candidates for ASD regulate genes that converge on the same key pathways.

**Figure 5.**
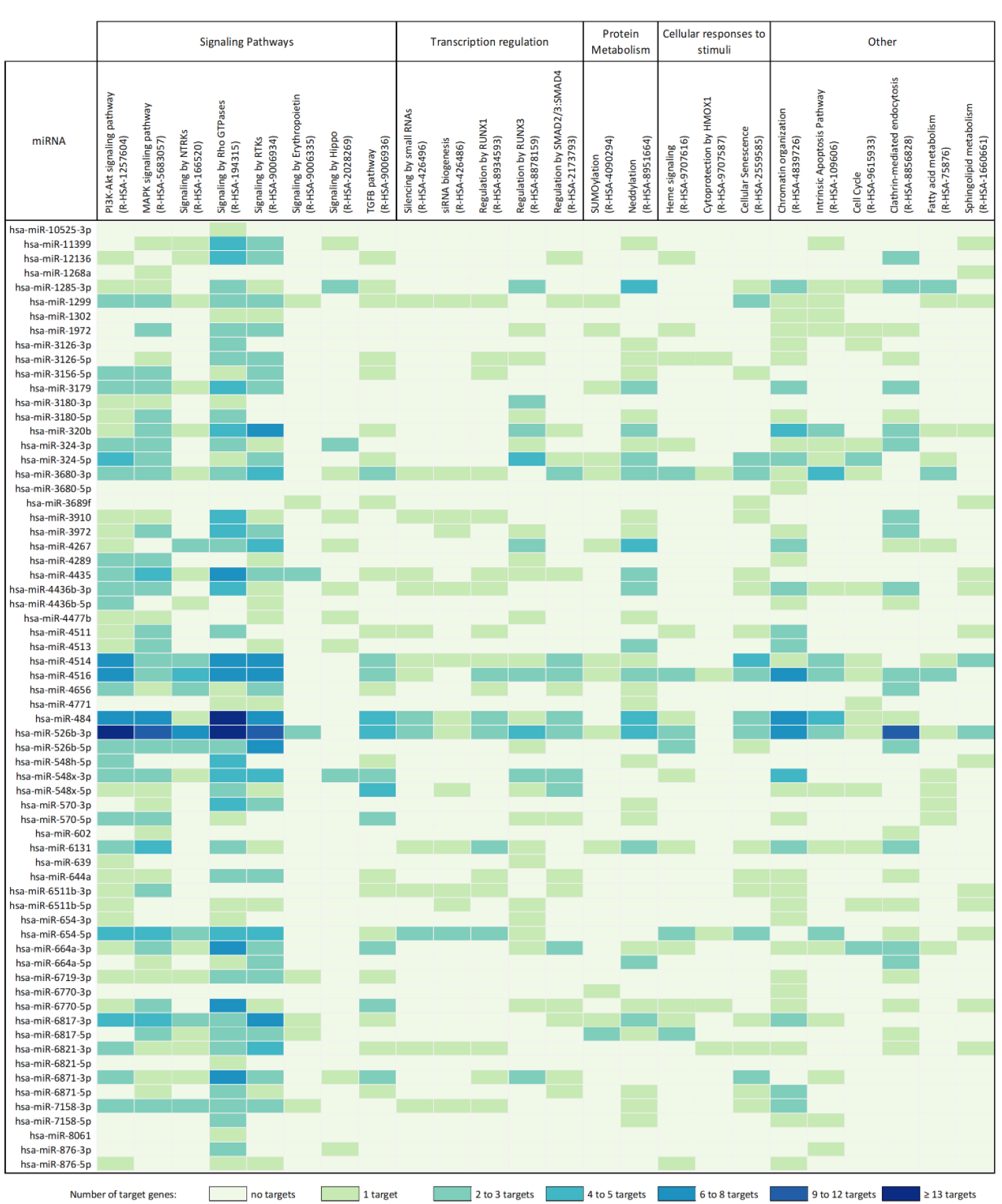
Schematic representation of miRNA regulatory pathways identified through functional enrichment analysis of their target genes. Only statistically significantly enriched Reactome pathways (adjusted *p*-value < 0.05) are represented (see methods for detail). The colors represent the number of miRNA target genes enriched in each pathway (darker colors are for high numbers of target genes, while lighter colors are for low numbers of target genes). The miRNAs that do not target a gene in the represented pathways are not shown is this scheme. **Legend:** NTRKs, Neurotrophin Tyrosine Kinase Receptors; RTKs, Receptor Tyrosine Kinases; TGFB, transforming growth factor-beta.

## Discussion

The high heritability of ASD has been recognized for decades, and hundreds of genes were previously associated to the disorder, yet the genetic determinants remain unknown for most patients (8–11). This is partly due to the scarcity of studies that comprehensively examine the consequences of variation in the entire genome, with non-coding regions and regulatory mechanisms remaining insufficiently explored. A clear example is miRNAs genes, which are frequently located in intergenic regions or within introns. Although miRNAs are key post-transcriptional regulators essential for brain development and function, the impact of variants within these genes remains poorly understood.

Seeking to advance knowledge of regulatory gene mechanisms in ASD, in this study we leveraged large genomic datasets for a detailed examination of miRNA gene variants in ASD patients and control subjects. We report a comprehensive gene burden analysis aggregating the effects of multiple genetic variants within each miRNA gene into a single statistic for each gene. Two complementary strategies, targeting SNV and CNV identification, were employed to detect rare variants predicted to affect miRNA structure and/or function. Our analysis specifically examined variants encoding the hairpin structural region, which are essential for correct miRNA processing, stability and target specificity.

This work led to the discovery of 28 miRNA genes significantly enriched for putative disruptive SNVs in ASD cases and 30 miRNA genes exclusively or more frequently targeted by CNVs in ASD individuals when compared to controls. The variants selected in these miRNA genes are predicted to alter the expression levels of mature miRNAs and/or disturb the recognition between mature miRNAs and miRNA targets, in turn impacting the expression of miRNA target genes. Overall, these 58 miRNA genes encode 70 mature miRNAs that are strong candidates for involvement in ASD. Of these, 49 mature miRNAs represent novel ASD candidates, which in some cases, namely hsa-miR-3179, hsa-miR-3180, hsa-miR-4435 and hsa-miR-4771, had already been associated with Schizophrenia, a disorder that phenotypically and genetically overlaps with ASD (19). The results also provide supportive evidence for 21 previously implicated mature miRNAs. From these, hsa-miR-324-5p, hsa-miR-484 and hsa-miR-4516 were already associated with ASD in both genetic and gene expression studies underscoring their likely important role in the disorder (17,21,77,84,87). An additional 15 mature miRNAs were reported to be deleted or duplicated in previous CNV studies (84,85,87), while 6 were identified in at least two gene expression studies (23,24,68,74,88,89).

Examination of the gene targets for the identified miRNAs show that most are expressed in the adult brain as well as during critical stages of brain development, suggesting a role of these miRNAs in essential brain processes. In line with this, gene ontology analysis showed that the miRNA targets are enriched in synaptic structures and regulatory complexes, further supporting their involvement in brain processes. Functional analysis identified shared biological pathways among miRNA targets, including genes involved in neuronal development, signaling transduction (notably the MAPK, PI3K-Akt, Neurotrophin, and RHO GTPase signaling pathways), chromatin modification, protein metabolism, and transcription regulation. All these pathways have been implicated in ASD (6,7,9,11,17), reinforcing the importance of these miRNAs in the regulation of a diversity of ASD-associated gene networks.

An important attribute of miRNAs is their ability to regulate multiple genes while, conversely, individual genes can be targeted by different miRNAs. This complex and interconnected nature of post-transcriptional regulation was also clearly evident from the network analysis performed in this study. In this analysis, we found that 44 of the candidate mature miRNAs are predicted to regulate 71 high-confidence ASD genes from the SFARI database, with individual miRNAs regulating up to 9 different ASD candidate genes. The genes categorized as high confidence in the carefully curated SFARI database meet the strongest evidence criteria from multiple large-scale studies, including replication across multiple cohorts, robust statistical association, and/or functional evidence for a role in brain development or function. We also found evidence that strong candidate genes for ASD can be regulated by multiple miRNAs, for instance the strongly ASD-associated genes KCNB1, MECP2, NCKAP1 and ZBTB20 are each regulated by at least 4 miRNAs identified in this study. This apparent bi-directional regulatory redundancy ensures that a robust control of key pathways is in place. Taken together, the diverse potentially disruptive variants identified in miRNA genes in ASD patients converge on common pathways and biological functions. This not only reinforces their shared involvement in ASD pathophysiology, but provides strong evidence for a central role of miRNA mediated regulation in this disorder.

The overall design of this study integrated strategies that ensure reliable results and a meaningful contribution to advancing the understanding of ASD. On one hand, SNVs were identified using genetic data from a large cohort with WGS from 4300 ASD patients. This approach allowed the detection of more variants in miRNA genes than previous studies, since most of the miRNA genes are located outside protein-coding regions and thus are not covered by exome arrays or whole exome-sequencing. This work also went beyond other studies by including additional functional miRNA gene regions for analysis, namely not only variants located on seed and mature regions, but also in the basal and apical motifs of the miRNA sequence, or predicted to change the structure of the hairpin. Furthermore, the CNV datasets were re-annotated with recent information from miRNA databases, which allowed the re-analysis of CNVs that were previously considered non-genic, leading to the discovery of new miRNAs in ASD patients. Indeed, while we identified miRNAs that were described in previous ASD genetic studies, we also found multiple miRNAs that have only recently been annotated by MIRBASE v22, and therefore could not be detected by older studies. Finally, the functional impact of these miRNAs was evaluated through examination of regulatory networks and enrichment analysis of miRNA target genes, highlighting shared biological functions and pathways previously implicated in ASD (6,7,9,11,17). The overall result was a complex, convergent regulatory network, reflecting the complexity of this brain dysfunction and underscoring the need for regulatory safeguards in which miRNAs likely play a crucial role.

Future research should experimentally validate the predicted miRNA–target interactions identified in this study, and evaluate their impact on target gene expression levels. Our analysis did not correlate our findings with clinical data, and bridging the gene-pathways-phenotype diversity gap will be essential to advance our understanding of ASD and to translate biological insights into clinical applications. Given that many of the enriched pathways (e.g., MAPK, PI3K–Akt, and chromatin modification) overlap, investigating their crosstalk in ASD and other neurodevelopmental disorders could pave the way for biomarker discovery and development of targeted therapies.

In conclusion, our findings provide further robust evidence for the involvement of miRNAs in the neurodevelopmental pathways, synaptic structures, and signaling mechanisms underlying ASD, emphasizing the complex interplay of molecular factors that contribute to the disorder. Of note, the discovery of potentially disruptive variants in miRNAs previously shown to be altered in gene expression or miRNA profiling studies of ASD patients reinforces their relevance for diagnostic and risk assessment. Future work integrating genetic, miRNA expression, and comprehensive clinical data, including signs and symptoms that go beyond neurodevelopment and behavior, which may be of interest in terms of research, will be crucial for a deeper understanding of the complex networks underlying ASD and to enable translation into clinical practice.

## Acknowledgments

The authors would like to acknowledge the resources of MSSNG (www.mss.ng, Autism Speaks and The Centre for Applied Genomics at The Hospital for Sick Children, Toronto, Canada) the Autism Genome Project (AGP) and the Simons Foundation Autism Research Initiative (SFARI) Simplex Collection (SSC) for facilitating access to the datasets used for the analysis described in this manuscript. The datasets were obtained from MSSNG resource (www.mss.ng), the study Sanders et al., 2011 and dbGaP through the accession number phs000267.v5.p2. We also thank the participating families for their time and contributions to these databases, as well as the generosity of the donors who supported these programs.

This research was supported by Fundação para a Ciência e a Tecnologia (UID/04046/2023 to Instituto de Biosistemas & Ciências Integrativas and UID/00006/2025 to Centro de Estatística e Aplicações (DOI: 10.54499/UIDP/00006/2020)), by PAC-POCI-01-0145-FEDER-016428 MEDPERSYST, by

DeePer—Deep graph learning approaches to personalized medicine (EXPL/CCI-BIO/0126/2021), and by National Institute of Health Doutor Ricardo Jorge. M.A., A.R.M. and J.V. were the recipients of BioSys PhD programme fellowship from FCT (Portugal) with references PD/BD/52485/2014, PD/BD/113773/2015, and PD/BD/131390/2017, respectively.

## Author contributions

A.R.M and A.M.V. design the study. A.R.M., H.M., L.S. and A.M.V. developed the methodology. A.R.M., M.A. and J.V. conducted the investigation. A.R.M. and H.M. performed data curation. A.R.M. performed data visualization and validation. A.M.V. and G.O. provided resources and acquired funding. A.M.V. supervised the study. A.R.M. and A.M.V. wrote the original manuscript draft and all authors critically reviewed and edited the manuscript.

## Supplement Description

Supplement Methods, Results, Figures S1-S2, Table S1-S9 (excel file)

Supplement Description: Supplement Methods, Results, Figures S1-S9, Tables S1-S4

## References

1. Zeidan J, Fombonne E, Scorah J, Ibrahim A, Durkin MS, Saxena S, et al. (2022): Global prevalence of autism: A systematic review update. Autism Res 15: 778–790.

2. American Psychiatric Association (2013): Diagnostic and Statistical Manual of Mental Disorders: DSM-5, 5th ed. American Psychiatric Association Publishing. 10.1176/appi.books.9780890425596

3. Bai D, Yip BHK, Windham GC, Sourander A, Francis R, Yoffe R, et al. (2019): Association of Genetic and Environmental Factors With Autism in a 5-Country Cohort. JAMA Psychiatry 76: 1035–1043.

4. Sandin S, Yip BHK, Yin W, Weiss LA, Dougherty JD, Fass S, et al. (2024): Examining Sex Differences in Autism Heritability. JAMA Psychiatry 81: 673–680.

5. Sanders SJ, Ercan-Sencicek AG, Hus V, Luo R, Murtha MT, Moreno-De-Luca D, et al. (2011): Multiple recurrent de novo CNVs, including duplications of the 7q11.23 Williams syndrome region, are strongly associated with autism. Neuron 70: 863–885.

6. Pinto D, Delaby E, Merico D, Barbosa M, Merikangas A, Klei L, et al. (2014): Convergence of genes and cellular pathways dysregulated in autism spectrum disorders. Am J Hum Genet 94: 677–694.

7. De Rubeis S, He X, Goldberg AP, Poultney CS, Samocha K, Cicek AE, et al. (2014): Synaptic, transcriptional and chromatin genes disrupted in autism. Nature 515: 209–215.

8. Satterstrom FK, Kosmicki JA, Wang J, Breen MS, De Rubeis S, An J-Y, et al. (2020): Large-Scale Exome Sequencing Study Implicates Both Developmental and Functional Changes in the Neurobiology of Autism. Cell 180: 568-584.e23.

9. Yuen RKC, Merico D, Bookman M, L Howe J, Thiruvahindrapuram B, Patel RV, et al. (2017): Whole genome sequencing resource identifies 18 new candidate genes for autism spectrum disorder. Nat Neurosci 20: 602–611.

10. Mahjani B, De Rubeis S, Gustavsson Mahjani C, Mulhern M, Xu X, Klei L, et al. (2021): Prevalence and phenotypic impact of rare potentially damaging variants in autism spectrum disorder. Mol Autism 12: 65.

11. Trost B, Thiruvahindrapuram B, Chan AJS, Engchuan W, Higginbotham EJ, Howe JL, et al. (2022): Genomic architecture of autism from comprehensive whole-genome sequence annotation. Cell 185: 4409-4427.e18.

12. Stott J, Wright T, Holmes J, Wilson J, Griffiths-Jones S, Foster D, Wright B (2023): A systematic review of non-coding RNA genes with differential expression profiles associated with autism spectrum disorders. PloS One 18: e0287131.

13. Friedman RC, Farh KK-H, Burge CB, Bartel DP (2009): Most mammalian mRNAs are conserved targets of microRNAs. Genome Res 19: 92–105.

14. Bartel DP (2018): Metazoan MicroRNAs. Cell 173: 20–51.

15. Gulyaeva LF, Kushlinskiy NE (2016): Regulatory mechanisms of microRNA expression. J Transl Med 14: 143.

16. Davis GM, Haas MA, Pocock R (2015): MicroRNAs: Not “Fine-Tuners” but Key Regulators of Neuronal Development and Function. Front Neurol 6: 245.

17. Wu YE, Parikshak NN, Belgard TG, Geschwind DH (2016): Genome-wide, integrative analysis implicates microRNA dysregulation in autism spectrum disorder. Nat Neurosci 19: 1463–1476.

18. Sánchez-Mora C, Soler Artigas M, Garcia-Martínez I, Pagerols M, Rovira P, Richarte V, et al. (2019): Epigenetic signature for attention-deficit/hyperactivity disorder: identification of miR-26b-5p, miR-185-5p, and miR-191-5p as potential biomarkers in peripheral blood mononuclear cells. Neuropsychopharmacol Off Publ Am Coll Neuropsychopharmacol 44: 890–897.

19. Warnica W, Merico D, Costain G, Alfred SE, Wei J, Marshall CR, et al. (2015): Copy Number Variable MicroRNAs in Schizophrenia and Their Neurodevelopmental Gene Targets. Biol Psychiatry 77: 158–166.

20. Abu-Elneel K, Liu T, Gazzaniga FS, Nishimura Y, Wall DP, Geschwind DH, et al. (2008): Heterogeneous dysregulation of microRNAs across the autism spectrum. Neurogenetics 9: 153–161.

21. Kalemaj Z, Marino MM, Santini AC, Tomaselli G, Auti A, Cagetti MG, et al. (2022): Salivary microRNA profiling dysregulation in autism spectrum disorder: A pilot study. Front Neurosci 16: 945278.

22. Wang Z, Lu T, Li X, Jiang M, Jia M, Liu J, et al. (2022): Altered Expression of Brain-specific Autism-Associated miRNAs in the Han Chinese Population. Front Genet 13: 865881.

23. Salloum-Asfar S, Elsayed AK, Elhag SF, Abdulla SA (2021): Circulating Non-Coding RNAs as a Signature of Autism Spectrum Disorder Symptomatology. Int J Mol Sci 22: 6549.

24. Nguyen LS, Lepleux M, Makhlouf M, Martin C, Fregeac J, Siquier-Pernet K, et al. (2016): Profiling olfactory stem cells from living patients identifies miRNAs relevant for autism pathophysiology. Mol Autism 7: 1.

25. Mor M, Nardone S, Sams DS, Elliott E (2015): Hypomethylation of miR-142 promoter and upregulation of microRNAs that target the oxytocin receptor gene in the autism prefrontal cortex. Mol Autism 6: 46.

26. Talebizadeh Z, Butler MG, Theodoro MF (2008): Feasibility and relevance of examining lymphoblastoid cell lines to study role of microRNAs in autism. Autism Res Off J Int Soc Autism Res 1: 240–250.

27. Ragusa M, Santagati M, Mirabella F, Lauretta G, Cirnigliaro M, Brex D, et al. (2020): Potential Associations Among Alteration of Salivary miRNAs, Saliva Microbiome Structure, and Cognitive Impairments in Autistic Children. Int J Mol Sci 21: 6203.

28. Rahnama M, Abdul-Tehrani H, Mohammadi MR, Mirzaie M, Jahandideh P, Memari A (2024): Expression analysis of microRNAs as candidate biomarkers in Iranian children with autism spectrum disorder. J Neurorestoratology 12: 100096.

29. Toma C, Torrico B, Hervás A, Salgado M, Rueda I, Valdés-Mas R, et al. (2015): Common and rare variants of microRNA genes in autism spectrum disorders. World J Biol Psychiatry Off J World Fed Soc Biol Psychiatry 16: 376–386.

30. Williams SM, An JY, Edson J, Watts M, Murigneux V, Whitehouse AJO, et al. (2019): An integrative analysis of non-coding regulatory DNA variations associated with autism spectrum disorder. Mol Psychiatry 24: 1707–1719.

31. Wong A, Zhou A, Cao X, Mahaganapathy V, Azaro M, Gwin C, et al. (2022): MicroRNA and MicroRNA-Target Variants Associated with Autism Spectrum Disorder and Related Disorders. Genes 13: 1329.

32. Cammaerts S, Strazisar M, De Rijk P, Del Favero J (2015): Genetic variants in microRNA genes: impact on microRNA expression, function, and disease. Front Genet 6. 10.3389/fgene.2015.00186

33. Machowska M, Galka-Marciniak P, Kozlowski P (2022): Consequences of genetic variants in miRNA genes. Comput Struct Biotechnol J 20: 6443–6457.

34. Lu J, Zhu Y, Williams S, Watts M, Tonta MA, Coleman HA, et al. (2020): Autism-associated miR-873 regulates ARID1B, SHANK3 and NRXN2 involved in neurodevelopment. Transl Psychiatry 10: 418.

35. Karczewski KJ, Francioli LC, Tiao G, Cummings BB, Alföldi J, Wang Q, et al. (2020): The mutational constraint spectrum quantified from variation in 141,456 humans. Nature 581: 434–443.

36. MacDonald JR, Ziman R, Yuen RKC, Feuk L, Scherer SW (2014): The Database of Genomic Variants: a curated collection of structural variation in the human genome. Nucleic Acids Res 42: D986–D992.

37. Cammaerts S, Strazisar M, Dierckx J, Del Favero J, De Rijk P (2016): miRVaS: a tool to predict the impact of genetic variants on miRNAs. Nucleic Acids Res 44: e23–e23.

38. Oak N, Ghosh R, Huang K, Wheeler DA, Ding L, Plon SE (2019): Framework for microRNA variant annotation and prioritization using human population and disease datasets. Hum Mutat 40: 73–89.

39. Cooper GM, Coe BP, Girirajan S, Rosenfeld JA, Vu TH, Baker C, et al. (2011): A copy number variation morbidity map of developmental delay. Nat Genet 43: 838–846.

40. Shaikh TH, Gai X, Perin JC, Glessner JT, Xie H, Murphy K, et al. (2009): High-resolution mapping and analysis of copy number variations in the human genome: A data resource for clinical and research applications. Genome Res 19: 1682–1690.

41. Guo MH, Plummer L, Chan Y-M, Hirschhorn JN, Lippincott MF (2018): Burden Testing of Rare Variants Identified through Exome Sequencing via Publicly Available Control Data. Am J Hum Genet 103: 522–534.

42. Sticht C, De La Torre C, Parveen A, Gretz N (2018): miRWalk: An online resource for prediction of microRNA binding sites. PloS One 13: e0206239.

43. Chen Y, Wang X (2020): miRDB: an online database for prediction of functional microRNA targets. Nucleic Acids Res 48: D127–D131.

44. Huang H-Y, Lin Y-C-D, Cui S, Huang Y, Tang Y, Xu J, et al. (2022): miRTarBase update 2022: an informative resource for experimentally validated miRNA–target interactions. Nucleic Acids Res 50: D222–D230.

45. Tweedie S, Braschi B, Gray K, Jones TEM, Seal RL, Yates B, Bruford EA (2021): Genenames.org: the HGNC and VGNC resources in 2021. Nucleic Acids Res 49: D939–D946.

46. Kolberg L, Raudvere U, Kuzmin I, Adler P, Vilo J, Peterson H (2023): g:Profiler—interoperable web service for functional enrichment analysis and gene identifier mapping (2023 update). Nucleic Acids Res 51: W207–W212.

47. Gillespie M, Jassal B, Stephan R, Milacic M, Rothfels K, Senff-Ribeiro A, et al. (2022): The reactome pathway knowledgebase 2022. Nucleic Acids Res 50: D687–D692.

48. Asif M, Martiniano HFMC, Marques AR, Santos JX, Vilela J, Rasga C, et al. (2020): Identification of biological mechanisms underlying a multidimensional ASD phenotype using machine learning. Transl Psychiatry 10: 43.

49. Shannon P, Markiel A, Ozier O, Baliga NS, Wang JT, Ramage D, et al. (2003): Cytoscape: A Software Environment for Integrated Models of Biomolecular Interaction Networks. Genome Res 13: 2498–2504.

50. GTEx Consortium (2020): The GTEx Consortium atlas of genetic regulatory effects across human tissues. Science 369: 1318–1330.

51. Sjöstedt E, Zhong W, Fagerberg L, Karlsson M, Mitsios N, Adori C, et al. (2020): An atlas of the protein-coding genes in the human, pig, and mouse brain. Science 367: eaay5947.

52. Kavakiotis I, Alexiou A, Tastsoglou S, Vlachos IS, Hatzigeorgiou AG (2022): DIANA-miTED: a microRNA tissue expression database. Nucleic Acids Res 50: D1055–D1061.

53. Miller JA, Ding S-L, Sunkin SM, Smith KA, Ng L, Szafer A, et al. (2014): Transcriptional landscape of the prenatal human brain. Nature 508: 199–206.

54. Lindsay SJ, Xu Y, Lisgo SN, Harkin LF, Copp AJ, Gerrelli D, et al. (2016): HDBR Expression: A Unique Resource for Global and Individual Gene Expression Studies during Early Human Brain Development. Front Neuroanat 10. 10.3389/fnana.2016.00086

55. Ghahramani Seno MM, Hu P, Gwadry FG, Pinto D, Marshall CR, Casallo G, Scherer SW (2011): Gene and miRNA expression profiles in autism spectrum disorders. Brain Res 1380: 85–97.

56. Ander BP, Barger N, Stamova B, Sharp FR, Schumann CM (2015): Atypical miRNA expression in temporal cortex associated with dysregulation of immune, cell cycle, and other pathways in autism spectrum disorders. Mol Autism 6: 37.

57. Huang F, Long Z, Chen Z, Li J, Hu Z, Qiu R, et al. (2015): Investigation of Gene Regulatory Networks Associated with Autism Spectrum Disorder Based on MiRNA Expression in China. PloS One 10: e0129052.

58. Mundalil Vasu M, Anitha A, Thanseem I, Suzuki K, Yamada K, Takahashi T, et al. (2014): Serum microRNA profiles in children with autism. Mol Autism 5: 40.

59. Hicks SD, Ignacio C, Gentile K, Middleton FA (2016): Salivary miRNA profiles identify children with autism spectrum disorder, correlate with adaptive behavior, and implicate ASD candidate genes involved in neurodevelopment. BMC Pediatr 16: 52.

60. Bleazard T (2018): Investigating the Role of microRNAs in Autism. United Kingdom: The University of Manchester. Retrieved August 16, 2025, from https://www.semanticscholar.org/paper/Investigating-the-role-of-microRNAs-in-autism-Bleazard/0596681fc0afbe76a331d74b995d89e5d4908f5e

61. Cirnigliaro M, Barbagallo C, Gulisano M, Domini CN, Barone R, Barbagallo D, et al. (2017): Expression and Regulatory Network Analysis of miR-140-3p, a New Potential Serum Biomarker for Autism Spectrum Disorder. Front Mol Neurosci 10: 250.

62. Kichukova TM, Popov NT, Ivanov IS, Vachev TI (2017): Profiling of Circulating Serum MicroRNAs in Children with Autism Spectrum Disorder using Stem-loop qRT-PCR Assay. Folia Med (Plovdiv) 59: 43–52.

63. Pagan C, Goubran-Botros H, Delorme R, Benabou M, Lemière N, Murray K, et al. (2017): Disruption of melatonin synthesis is associated with impaired 14-3-3 and miR-451 levels in patients with autism spectrum disorders. Sci Rep 7: 2096.

64. Popov N, Minchev D, Naydenov M, Minkov I, Vachev T (2018): Investigation of Circulating Serum MicroRNA-328-3p and MicroRNA-3135a Expression as Promising Novel Biomarkers for Autism Spectrum Disorder. Balk J Med Genet BJMG 21: 5–12.

65. Vaccaro T da S, Sorrentino JM, Salvador S, Veit T, Souza DO, de Almeida RF (2018): Alterations in the MicroRNA of the Blood of Autism Spectrum Disorder Patients: Effects on Epigenetic Regulation and Potential Biomarkers. Behav Sci Basel Switz 8. 10.3390/bs8080075

66. Yu D, Jiao X, Cao T, Huang F (2018): Serum miRNA expression profiling reveals miR-486-3p may play a significant role in the development of autism by targeting ARID1B. Neuroreport 29: 1431–1436.

67. Jyonouchi H, Geng L, Toruner GA, Rose S, Bennuri SC, Frye RE (2019): Serum microRNAs in ASD: Association With Monocyte Cytokine Profiles and Mitochondrial Respiration. Front Psychiatry 10: 614.

68. Nakata M, Kimura R, Funabiki Y, Awaya T, Murai T, Hagiwara M (2019): MicroRNA profiling in adults with high-functioning autism spectrum disorder. Mol Brain 12: 82.

69. Ozkul Y, Taheri S, Bayram KK, Sener EF, Mehmetbeyoglu E, Öztop DB, et al. (2020): A heritable profile of six miRNAs in autistic patients and mouse models. Sci Rep 10: 9011.

70. Sehovic E, Spahic L, Smajlovic-Skenderagic L, Pistoljevic N, Dzanko E, Hajdarpasic A (2020): Identification of developmental disorders including autism spectrum disorder using salivary miRNAs in children from Bosnia and Herzegovina. PloS One 15: e0232351.

71. Hicks SD, Carpenter RL, Wagner KE, Pauley R, Barros M, Tierney-Aves C, et al. (2020): Saliva MicroRNA Differentiates Children With Autism From Peers With Typical and Atypical Development. J Am Acad Child Adolesc Psychiatry 59: 296–308.

72. Cui L, Du W, Xu N, Dong J, Xia B, Ma J, et al. (2021): Impact of microRNAs in interaction with environmental factors on autism spectrum disorder: an exploratory pilot study. Front Psychiatry 12: 715481.

73. Frye RE, Rose S, McCullough S, Bennuri SC, Porter-Gill PA, Dweep H, Gill PS (2021): MicroRNA expression profiles in autism spectrum disorder: role for miR-181 in immunomodulation. J Pers Med 11: 922.

74. Kichukova T, Petrov V, Popov N, Minchev D, Naimov S, Minkov I, Vachev T (2021): Identification of serum microRNA signatures associated with autism spectrum disorder as promising candidate biomarkers. Heliyon 7. Retrieved March 10, 2025, from https://www.cell.com/heliyon/fulltext/S2405-8440(21)01565-6

75. Zhang Y, Pang Y, Feng W, Jin Y, Chen S, Ding S, et al. (2022): miR-124 regulates early isolation-induced social abnormalities via inhibiting myelinogenesis in the medial prefrontal cortex. Cell Mol Life Sci CMLS 79: 507.

76. Hosokawa R, Yoshino Y, Funahashi Y, Horiuchi F, Iga J, Ueno S (2022): MiR-15b-5p expression in the peripheral blood: a potential diagnostic biomarker of autism spectrum disorder. Brain Sci 13: 27.

77. Gill PS, Dweep H, Rose S, Wickramasinghe PJ, Vyas KK, McCullough S, et al. (2022): Integrated microRNA– mRNA Expression Profiling Identifies Novel Targets and Networks Associated with Autism. J Pers Med 12: 920.

78. Elsheikh MS, Ashaat EA, Ramadan A, Mohamed NH, Elaraby NM, El-Hariri HM, et al. (2023): Efficacy of Laser Acupuncture for Children With Autism Spectrum Disorder: Clinical, Molecular, and Biochemical Study. Pediatr Neurol 147: 44–51.

79. Zhao S, Zhong Y, Shen F, Cheng X, Qing X, Liu J (2024): Comprehensive exosomal microRNA profile and construction of competing endogenous RNA network in autism spectrum disorder: A pilot study. Biomol Biomed 24: 292.

80. Karagöz H, Akça ÖF, Yildirim MS, Zamani AG, Oflaz MB (2024): Comparison of MicroRNA Levels of 18-60-month-old Autistic Children with Those of Their Siblings and Controls. Clin Psychopharmacol Neurosci 22: 322.

81. Ali SJT, Khalaj-Kondori M, Feizi MAH, Haghi M (2024): Expression Levels of miR-124a, miR-545-3p and BDNF in the Peripheral Blood Mononuclear Cells Are Associated with the Severity of Autism. Rep Biochem Mol Biol 13: 1.

82. Li Y, Liu C, Jin Q, Yu H, Long H (2025): H19/miR-484 axis serves as a candidate biomarker correlated with autism spectrum disorder. Int J Dev Neurosci 85: e10403.

83. Ren J, Bai Y, Gao J, Hou Y, Mao J, Gao F, Wang J (2025): Diagnostic Value of Serum miR-499a-5p in Chinese Children with Autism Spectrum Disorders. J Mol Neurosci 75: 1–9.

84. Vaishnavi V, Manikandan M, Tiwary BK, Munirajan AK (2013): Insights on the functional impact of microRNAs present in autism-associated copy number variants. PloS One 8: e56781.

85. Marrale M, Albanese NN, Calì F, Romano V (2014): Assessing the impact of copy number variants on miRNA genes in autism by Monte Carlo simulation. PloS One 9: e90947.

86. The Autism Spectrum Disorders Working Group of The Psychiatric Genomics Consortium (2017): Metaanalysis of GWAS of over 16,000 individuals with autism spectrum disorder highlights a novel locus at 10q24.32 and a significant overlap with schizophrenia. Mol Autism 8: 21.

87. Qiu S, Qiu Y, Li Y, Zhu X, Liu Y, Qiao Y, et al. (2023): Nexus between genome-wide copy number variations and autism spectrum disorder in Northeast Han Chinese population. BMC Psychiatry 23: 96.

88. Guiducci L, Cabiati M, Santocchi E, Prosperi M, Morales MA, Muratori F, et al. (2023): Expression of miRNAs in pre-schoolers with Autism Spectrum disorders compared with typically developing peers and its effects after Probiotic supplementation. J Clin Med 12: 7162.

89. Mordaunt CE, Park BY, Bakulski KM, Feinberg JI, Croen LA, Ladd-Acosta C, et al. (2019): A meta-analysis of two high-risk prospective cohort studies reveals autism-specific transcriptional changes to chromatin, autoimmune, and environmental response genes in umbilical cord blood. Mol Autism 10: 36.

